# MatClassRSA: A Matlab toolbox for M/EEG classification and visualization of proximity matrices

**DOI:** 10.1101/194563

**Authors:** Bernard C. Wang, Anthony M. Norcia, Blair Kaneshiro

## Abstract

MatClassRSA is a Matlab toolbox that performs M/EEG classification and produces various visualizations of the resulting confusion matrices. This toolbox is aimed toward cognitive neuroscience researchers who wish to apply classification-style analyses to repeated trials of data. The functionalities of the toolbox fall into three categories: (1) M/EEG classification, (2) computation of Representational Dissimilarity Matrices (RDMs) from confusion or other proximity matrices, and (3) clustering and tree visualizations of RDMs. MatClassRSA combines out-of-the-box functionality with a variety of customization options. Usage of the toolbox requires only a high-level understanding of machine learning and the ability to call Matlab functions from scripts.

## 1 Motivation and significance

Classification—the task of assigning categorical labels to observations of test data, based upon a statistical model built from labeled training observations [1]—has been applied to brain responses from various recording modalities including fMRI, ECoG, and M/EEG [2]. In this context, classifiers attempt to predict stimulus attributes, cognitive states, motor activations, and more from brain responses [3]. EEG classification has developed primarily in the context of Brain-Computer Interfaces (BCIs), where it is still widely used [4]. Here, the goal is to enable users who cannot communicate verbally or through other means (such as typing) to control a computer or other external apparatus through brain activity alone. An EEG-BCI can involve decoding of imagery, typically imagined movements [5], or selective attention to visual and auditory stimuli using steady-state evoked responses (SS-EPs) [6, 7] or event-related potentials (ERPs) [8, 9].

Successful BCIs require high decoding accuracy—that is, the system must be able to correctly label brain data, typically on a single-trial basis. In recent years, however, M/EEG classification has extended beyond BCI into the realm of cognitive neuroscience, and has been used to study representation of object [10, 11] and language [12] categories, as well as performance on cognitive tasks [13]. Some studies have identified relevant spatiotemporal features of cortical responses by interpreting classifier weights [11, 14] or by employing a ‘searchlight’ approach [15, 10]. In contrast to BCI applications, here the fine-tuning of classification algorithms for slight performance gains is not the main goal; however, a properly functioning classifier is necessary in order to produce valid and interpretable results.

More recently, M/EEG classification has also proved useful in the context of Representational Similarity Analysis (RSA). This approach enables the structure of a set of stimuli to be compared across imaging, computational, and behavioral modalities [16] by abstracting raw responses into pairwise distances which are summarized in square, symmetric matrices termed Representational Dissimilarity Matrices (RDMs). In RSA studies using fMRI, RDMs have been computed by correlating multivariate voxel data for stimulus pairs [16, 17]. M/EEG RDMs have additionally been computed using pairwise classification accuracies [18, 19] and multicategory confusion matrices [10]. Recent studies demonstrate the value of M/EEG classification in cognitive neuroscience research. Moreover, researchers who perform averaging-based ERP analyses may already have classification-ready M/EEG data, as collections of short, repeated trials reflecting various categories are generally amenable to classification analyses. However, the domain-specific expertise needed to implement classifiers and visualization tools may slow adoption among this research community. Therefore, M/EEG researchers could benefit from easy-to-use software that performs classifications and creates RSA-style visualizations of the classifier output. These are the contributions of MatClassRSA. The toolbox handles any input M/EEG data that can be reshaped into a trials-by-feature matrix, where features can be time-sampled voltages, time-frequency coefficients, or spectral coefficients, and can be sampled from single or multiple electrodes. The present software provides a means for classifying and visualizing the data, and for ease of use is compatible with data matrices of the format output by the Matlab EEGLAB toolbox [20].

There exist a number of open-source software packages that provide complementary functionalities to MatClassRSA. Matlab implementations include BCILAB [21], which provides classification (offline and online) functionalities for BCI. MatClassRSA differs on the basis of having a cognitive neuroscience focus and added RSA visualization capabilities. The PRoNTo toolbox [22] also performs classification and regression analyses, but in its current implementation is focused on PET/fMRI imaging (NIfTI) input files rather than M/EEG data matrices. The MANIA [23] and MVPA^1^ toolboxes are similarly positioned toward imaging data. The RSA toolbox [24] performs a number of analyses including computation of RDMs, comparison of RDMs, and statistical analyses to quantify relatedness among RDMs. The toolbox appears to be developed more for fMRI and cell recordings than for M/EEG; however, classification-based RDMs computed using MatClassRSA can be subsequently input to the RSA toolbox for quantitative comparison with other brain and model RDMs. Finally, the DDTBOX performs EEG classification and related statistical analyses [25]. MatClassRSA provides complementary classifier options and visualization capabilities to this toolbox.

M/EEG preprocessing and machine learning libraries are also implemented in Python, including MNE [26, 27], PyMVPA [28], and scikit-learn [29]. However, for M/EEG researchers working primarily or solely in Matlab, a transition to Python presents a steeper learning curve and potentially less compatibility with existing data-analysis workflows. The present toolbox therefore enables researchers to remain in a familiar Matlab environment, with consistent formats for analysis scripts, variables, and data files.

The remainder of this paper is structured as follows. In Section 2 we provide an overview of the software architecture and its main functions. Illustrative examples are provided in Section 3, and we conclude with a discussion of impact and planned future developments of the software in Section 4.

## 2 Software description

The MatClassRSA toolbox is implemented in Matlab, and is verified to be compatible with versions R2016b and later, on Mac and Linux operating systems. The `libsvm` library [30] is required in order to use the SVM classifier.^2^ Users of this toolbox should have proficiency in Matlab at the level of calling scripts without the aid of a GUI, as this software is run by calling functions from a script or from the Matlab console. We assume that data have already been cleaned and epoched, and are stored in one of the acceptable data shapes described below. Input and output data are stored as variables; loading and saving data from and to .mat files is the responsibility of the user. While a high-level understanding of machine learning will be helpful, implementation expertise is not required. The example scripts provided with the toolbox can be extended to the analysis of new datasets and directory structures with minimal modifications.

### 2.1 Software Architecture and Functionalities

MatClassRSA comprises three main functionalities which are accessed through six user-invoked functions (Figure 1). First, M/EEG preparation and classification is performed by the function `classifyEEG`. Next, the `computeDistanceMatrix` function converts the confusion matrix output by the classifier to an RDM. The remaining four functions visualize RDMs: `plotMatrix` creates an image of a confusion matrix or RDM; `plotMDS` and `plotDendrogram` create clustering and hi-erarchical clustering plots, respectively, of an RDM, similar to those presented in previous RSA studies [17, 10]; and `plotMST` visualizes the RDM as a minimum spanning tree. Detailed descriptions of the inputs, outputs, and default parameters of the functions are included in the manual accompanying the toolbox.

**Figure 1:**
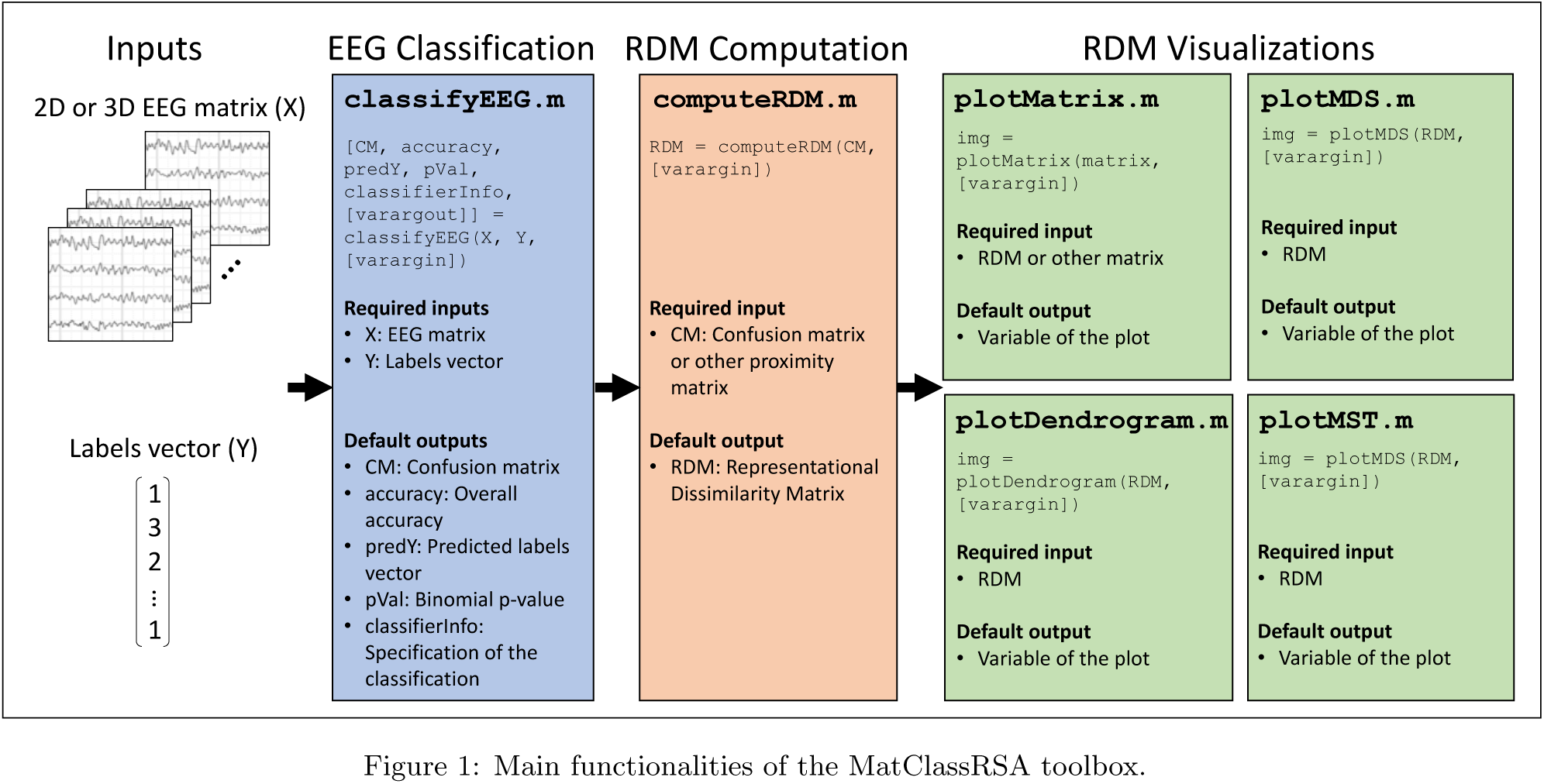
Main functionalities of the MatClassRSA toolbox.

#### 2.1.1 M/EEG Classification

The `classifyEEG` function requires two inputs: An M/EEG data frame X and a vector of stimulus labels Y. X can be a 3D space-by-feature-by-trial matrix, or a 2D trial-by-feature matrix. Y must correspond in length to the trial dimension of X.

The function accepts several optional data preparation and classification parameters. First, the user can specify whether the classification should be performed over a subset of data features (selected time points and/or electrodes). Researchers sometimes classify group-averaged rather than single trials [12, 31], and that option is provided here as well.

Cross validation is a common procedure for estimating prediction error [1]. Here, the collection of observations is partitioned into *K* roughly equally sized subsets, and each subset is used for testing once, while the other subsets are used to train the model in that ‘fold’ (generalizing to *K*-fold cross validation). MatClassRSA allows users to specify the number of folds *K* for cross validation, ranging from *K* = 2 to *K* = *N*, where *N* is the number of observations (leave-one-out cross validation). There is also an option to shuffle the ordering of the trials (while keeping them paired with the correct labels) prior to partitioning for cross validation.

Feature dimensionality reduction using PCA has been proposed for M/EEG classification [10], and the `classifyEEG` function allows users to specify a target number of PCs or desired proportion of variance explained for PCA decomposition along the feature dimension. If selected, PCA can be performed over the full collection of input data prior to cross validation (faster), or over just the training observations for each cross-validation fold (avoids the use of test data in constructing the classifier).

Three popular classifiers are supported: Support vector machine (SVM) [32], random forest (RF) [33], and linear discriminant analysis (LDA) [34], each with optional parameters that can be set by the user. The classification script has capabilities to assess statistical significance under the null distribution of the binomial distribution [10] or with permutation testing [35]. Finally, the user can specify a seed type for the random number generator that is used to shuffle trials and labels together, or just the labels during permutation tests.

The `classifyEEG` function outputs the confusion matrix, overall accuracy, vector of predicted labels, p-value, and a data struct containing the specifications of the classification. If permutation testing was performed, a vector of accuracies from each permutation iteration will additionally be returned. Finally, a *verbose* option can be invoked in the function call to cause the function to return the classifier models themselves.

#### 2.1.2 Computing Distance Matrices

The confusion matrix output by classifyEEG summarizes the performance of the classifier. With rows representing actual categories and columns predicted categories, element *CM*_*ij*_ of confusion matrix *CM* denotes the number of observations from category *i* that were labeled as being from category *j*. Values on the diagonal represent correct classification, and the sum of all elements in the matrix represents the total number of observations classified.

Multiclass confusion matrices can be treated as proximity matrices, with confusions serving as similarity measures [36]. Adapting the approach taken in a recent M/EEG classification study [10], `computeDistanceMatrix` is a generalized function for creating distance matrices of the type that are visualized for RSA. The function accepts as input any square matrix whose elements contain pairwise measures of similarity or distance (e.g., multiclass confusion matrices, correlation matrices, pairwise similarity or distance judgments), which we henceforth refer to as ‘proximity matrices’. The input proximity matrix can then be normalized (e.g., to produce self-similarity of 1 along the diagonal [37] or scale by the sum of the row, producing an estimated conditional probability matrix [38]) and symmetrized (arithmetic, geometric, or harmonic mean of the matrix and its transpose [37]), and values can be converted from similarities to distances (linear, power, or logarithmic). Finally, as some studies have operated not directly on distances but rather on percentile or ranked distances [16, 17], this functionality is also provided in the distance matrix computation.

The ability to customize or forgo any of the computations listed above enables users to input a variety of proximity matrices to the function. Users with precomputed RDMs can skip the `computeDistanceMatrix` function altogether and proceed directly to the visualization scripts; or, the function can be used to compute an RDM, which can then be analyzed further using the RSA toolbox [24].

#### 2.1.3 Visualizing Matrices

MatClassRSA provides four visualization options. First, `plotMatrix` images an input matrix (e.g., confusion matrix, proximity matrix, distance matrix). The function provides options for labeling rows and columns of the matrix with text or images, displaying text in the matrix cells, and colormap and colorbar display.

The second function, `plotMDS`, performs Multidimensional Scaling (MDS) to visualize the non-hierarchical and often high-dimensional structure of a distance matrix in two dimensions at a time [39]. This function includes options for setting the plot area, selecting MDS dimensions to be visualized, and labeling exemplars in the plot with text or images.

The hierarchical structure of a distance matrix can be visualized as a dendrogram using the `plotDendrogram` function. As the optimal choice of linkage may depend upon the type of distance (e.g., euclidean or non-euclidean) represented in a distance matrix [40], a variety of linkage options are available. The user can specify layout (e.g., orientation, leaf ordering, axis ranges) and labeling options for the plot.

Finally, a minimum spanning tree (MST) displays the shortest path among all objects in a set (i.e., classification categories), and can provide useful and complementary insights into the relationships among clusters in the data [41, 40]. Basic functionalities for generating minimum spanning trees are provided in the `plotMST` function. Here, the user may optionally specify text or image labels for the nodes of the tree.

#### 2.1.4 Code Utilization for Specific Use Cases

The MatClassRSA toolbox was designed to provide a general framework for M/EEG classification. As a result, certain more specific functionalities—such as single-electrode classification and pairwise classification—are not provided in the main functions. However, the software can be utilized to implement such use cases, and we provide example scripts that do so as part of the illustrative analyses in Section 3.

## 3 Illustrative Examples

We classify and visualize a publicly available collection of EEG data [42] that were published as part of a recent classification study [10]. In this study, each of 10 participants viewed a set of 72 images from 6 object categories 72 times (5,184 trials total; 72 trials per exemplar, 864 trials per category). The present analyses use data from participant 6 (S6.mat), who we found to produce the highest classification rates overall. We use the 3D EEG matrix X 3D as data matrix X and category-level labels vector `categoryLabels` as labels vector Y—a six-class classification problem—for the following analyses.

In the script `classificationExample.m`, we first classify single EEG trials. For replicability of these results, we set the ‘`randomSeed`’ parameter in the `classifyEEG` function call to ‘default’. We use default values for the remaining parameters of the function call: The ordering of trials is shuffled (but kept intact with respective labels) prior to cross validation; PCA is performed separately on the training partition in each cross-validation fold, retaining as many PCs needed to explain 90% of the variance along the feature dimension; and classification is performed with 10-fold cross validation using SVM with Gaussian (rbf) kernel. We use the binomial cumulative distribution function to estimate the *p*-value. This analysis produces an accuracy of 46.12% (binomial *p* < 0.001), similar to the results of the original published study [10]. We then input the resulting confusion matrix into the `computeDistanceMatrix` function, using default parameters: Each row of the input matrix is normalized by its respective diagonal entry; the matrix is then symmetrized based on the arithmetic mean of the matrix and its transpose; and similarities are converted to distances by *D* = 1 − *S*, with no ranking of distances.

Finally, we call each of the four visualization functions. We use the function `plotMatrix` to visualize the RDM. By default, each square on the matrix plot will be populated with a label giving the distance between the two corresponding classes on the x and y axes ticks. The ‘colormap’ parameter is set to ‘`summer`’ in order for the black distance labels to be visible. To set class labels on the x and y axes, we pass a cell array of text labels, {`‘HB’ ‘HF’ ‘AB’ ‘AF’ ‘FV’ ‘IO’`}, into the `‘axisLabels’` parameter. Then, we color the class labels with the `‘axisColors’` parameter by passing in the cell array {`[1 .5 .3] [.1 .5 1] ‘r’ ‘g’ ‘c’ ‘m’`} (RGB triplets and Matlab color abbreviations are among the acceptable color specifications). So that class labels and class colors are consistent for all visualizations, we use the same label and colors inputs in the calls to `plotMDS`, `plotDendrogram`, and `plotMST`, with the exception that the parameter names are changed to `‘nodeLabels’` and `‘nodeColors’` for these latter cases. The distance matrix, plot of MDS dimensions 1 and 2, dendrogram, and MST for this first classification are shown in Panel A of Figure 2.

**Figure 2:**
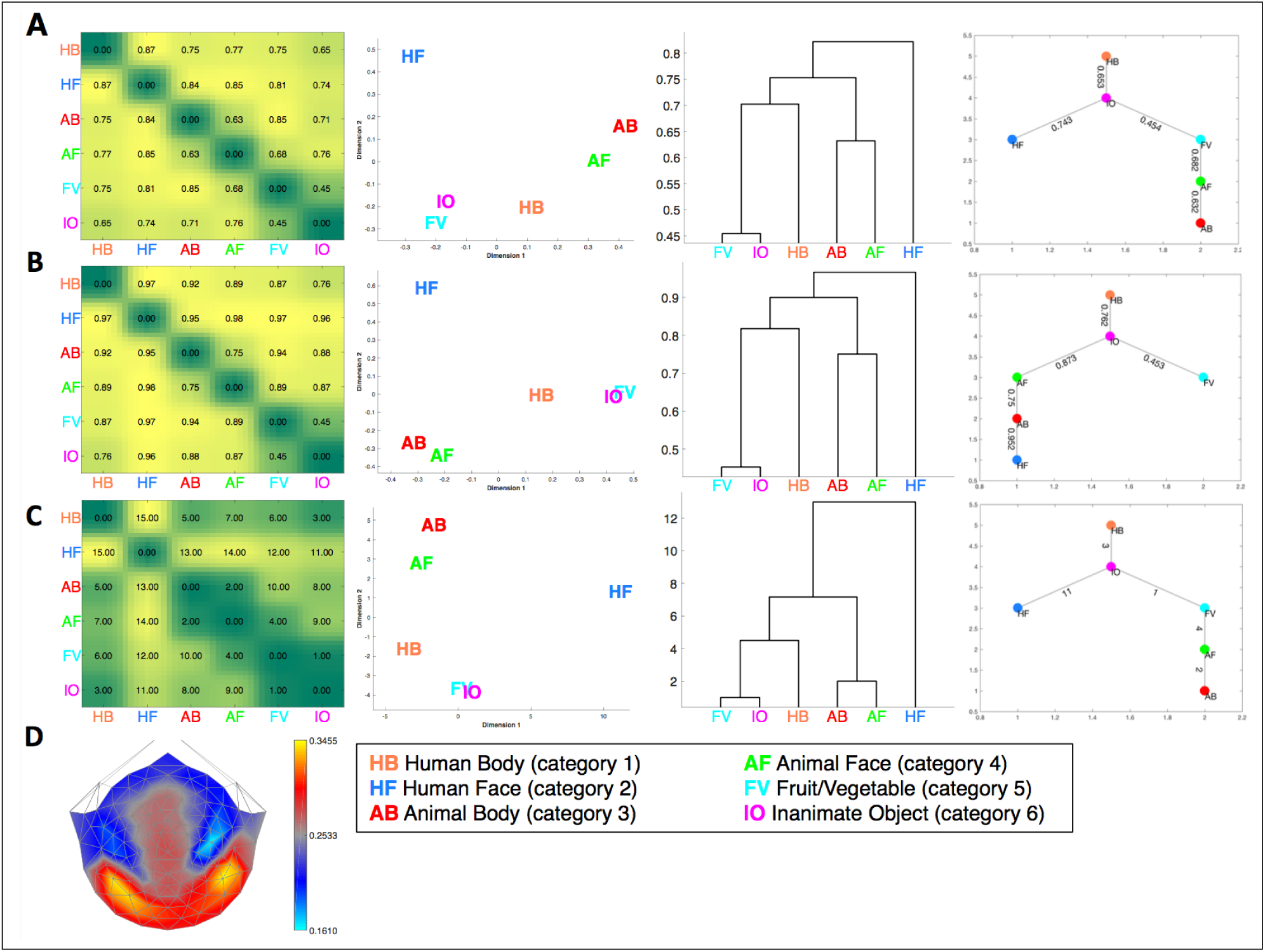
Results from illustrative analyses. A: Six-class single-trial classification results. B: Six-class classification results when trials are averaged in groups of five. C: Visualizations of distance matrix constructed from pairwise single-trial classification accuracies. D: Topographic plot of single-electrode classification rates for six-class single-trial classification.

As we have a fairly large number of trials for the six-class classification task, we repeat the above classification, now with trials from the same category averaged in groups of five. Keeping all other specifications of the classification the same as before, our accuracy now improves to 64.28% (binomial *p* < 0.001). The corresponding visualizations are shown in Panel B of Figure 2.

The script `classifyPairs.m` implements pairwise category-level classification of single EEG trials and visualization of the resulting collection of pairwise accuracies, an approach that has been used in a recent MEG-RSA study [19]. Here, we input to the `classifyEEG` function observations and labels from only two categories at a time. Rather than constructing the proximity matrix from the classifier confusions, we now construct it from each pairwise classification accuracy (higher accuracy implies greater distance between the two categories), and utilize only the rank-distance functionality of `computeDistanceMatrix`. The resulting plots are shown in Panel C of Figure 2, revealing a similar category structure to that produced by the previous multicategory classifications.

Finally, we consider classification of single electrodes. This ‘searchlight’-style analysis reveals which electrodes recorded data that are well discriminated by the classifier [15, 10]. In `classifySingleElectrodes.m`, we implement a loop to classify single-trial data from one electrode at a time (124 separate classifications) using the same classifier settings described above. The resulting single-electrode classification accuracies are visualized on a scalp map in Panel D of Figure 2. The helper functions used to produce the topoplot are included with the toolbox. Again, we find the results presented here to be broadly consistent with the previous published study [10].

### 4 Impact and Conclusions

We have introduced MatClassRSA, a Matlab toolbox for M/EEG classification and proximity matrix visualization. Classification is a powerful tool for studying M/EEG responses, and can extend the analysis of evoked responses to RSA-style analyses, permitting the study of large stimulus sets, comparison with other, disparate forms of data, and quantitative assessment of the underlying structure of the stimulus space. Through ‘searchlight’ approaches or analysis of classifier features, classification additionally provides a data-driven approach toward identifying features of the brain response that differentiate stimuli, tasks, or cognitive states. Finally, the nature of classification itself—assigning correct labels to unseen observations—points to potential clinical applications, where predictive decisions are key. While the MatClassRSA toolbox does not currently offer all of these potential functionalities, it does provide researchers the means to begin classifying and visualizing M/EEG data.

The current implementation of the toolbox enables researchers to perform basic classification analyses. MatClassRSA has been tested by M/EEG researchers from a number of research groups, and earlier versions of the code have been used for analyses that have been presented at scientific meetings. Development of this codebase is ongoing, and there are several avenues for further development. For example, other, more sophisticated classifiers, model and feature selection, and cross-validation procedures could be implemented. Additional statistical testing procedures, including multiple comparison correction, would be of value given the high dimensionality—in both time and space—of much M/EEG data. Visualization of results, too, could be extended beyond current offerings. It is hoped that the open-source publication of this software will encourage further developments and contributions from the research community.

In conclusion, MatClassRSA facilitates research in M/EEG classification, an area of growing interest in the field of cognitive neuroscience. Ease of use is provided with few function calls and out-of-the-box default settings for most forms of input data, while numerous input options to the main functions provide flexibility for more customized analyses. It is hoped that this toolbox will enable M/EEG researchers to approach their data in a new way through classification, providing an opportunity to explore multivariate analyses, RSA, and decoding paradigms.

## Footnotes

1 https://github.com/princetonuniversity/princeton-mvpa-toolbox

2 https://www.csie.ntu.edu.tw/~cjlin/libsvm/

## Acknowledgements

This research was supported by EY018875-04 (AMN), the Patrick Suppes Gift Fund (BK), and the Roberta Bowman Denning Fund for Humanities and Technology (BK). We gratefully acknowledge Yiran Duan, Steven Losorelli, and Duc T. Nguyen for testing earlier versions of the code; and Feng Ruan and Peter J. Kohler for helpful advice on implementations.

## Required Metadata

### Current code version

Ancillary data table required for subversion of the codebase. Kindly replace examples in right column with the correct information about your current code, and leave the left column as it is.

**Table 1:**
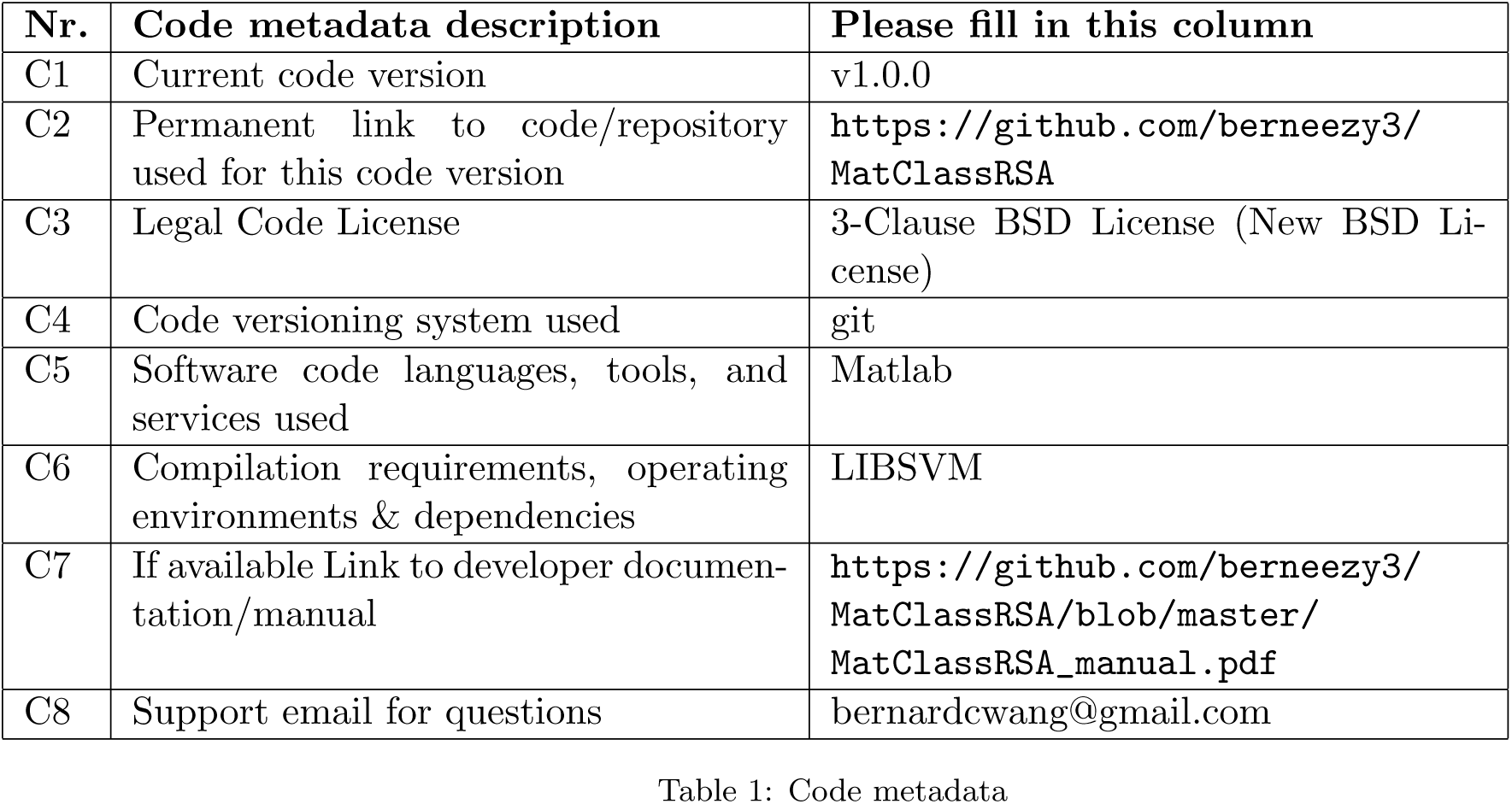
Code metadata

